# Unusual migratory strategy a key factor driving interactions at wind energy facilities in at-risk bats

**DOI:** 10.1101/2024.01.28.577637

**Authors:** Caitlin J. Campbell, David M. Nelson, Juliet Nagel, Jeff Clerc, Theodore J. Weller, Jamin G. Weiringa, Erin Fraser, Fred J. Longstaffe, Amanda M. Hale, Meghan Lout, Lori Pruitt, Robert Guralnick, Hannah B. Vander Zanden

## Abstract

Seasonal movement strategies are poorly understood for most animals, impeding broader understanding of processes underlying migration and limiting practical conservation needs. Here we develop and implement a framework for integrating multiple sources of endogenous markers, in particular stable hydrogen isotope data, that capture and scale dynamics from the movements of individuals to that of continental migration. We assembled and integrated thousands of new isotopic measurements from bat fur with existing datasets and applied this framework to reveal migratory patterns of three broadly distributed bat species most at risk for fatalities at wind energy facilities. Our findings show that the species comprising the lowest proportion of wind turbine fatalities (silver-haired bats) exhibits expected movements to lower latitudes in autumn and higher latitudes in spring. Surprisingly, the two species with higher wind turbine fatality rates (hoary and eastern red bats) have more complex movements, including significant movement to higher latitudes during autumn. We term this unique strategy “pell-mell” migration, during which some individuals are as likely to move to higher latitudes as lower latitudes, relative to their individual summering grounds, in early autumn, after which they move to similar or lower latitudes to overwinter. The pell-mell migratory period corresponds with peak fatalities at wind energy facilities, and bats moving northward during autumn are associated with mortality at those facilities. Our results provide direct support for the hypothesis that bat fatalities at wind energy facilities are related to migration and highlight the importance of migratory distance as an ultimate driver of increased interactions with wind energy facilities, which appears significantly greater for species that travel widely across latitudes in the autumn.

## Introduction

Animal migration is a spectacular natural phenomenon responsible for the movements of trillions of organisms, thousands of species, and kilotons of nutrients across the earth (1). Yet, we know shockingly little about the migratory strategies of most organisms because conventional techniques of studying migration are resource-intensive and limited to the study of large and easily-observed animals (2). Contrasting theories suggest that seasonal migration, here defined as a regular, repeated, and large-scale movement between different sites (3), is ultimately motivated by individuals tracking consistent climatic niches or resources (generally, food; [3–6]). However, proximate drivers governed by other social and behavioral factors, such as mating or ephemeral resource tracking, are not included in these general theories (8). Moving beyond coarse-scale understanding of migratory dynamics for less well-studied taxa remains a critical challenge in animal movement biology, and will require integrating complex individual-level behaviors and motivations to landscape-level shifts in the distributions of populations. New data types and analytical frameworks that scale from individuals to landscapes will both improve our understanding of animal migration and its drivers and likely uncover surprising and unique aspects of migration outside of well-studied examples.

Migratory tree-roosting bats (*sensu* [9]) are largely solitary, highly mobile, and notoriously difficult to study using conventional and even modern methods of migration biology (10, 11). Their impressive capacity for long-distance movement is well-known, but mostly anecdotal. For example, one species has colonized Hawaii, 3,665 km from continental North America; and three are regular vagrants in Bermuda, 1,064 km from mainland U.S. (12). Limited recapture data and location logging have shown long and often circuitous migratory routes of a few individuals (>1000 km, [12, 13]). Although migration theory predicts that niche or resource tracking across seasons would result in two discrete migratory periods between warm and cool seasons (15), tree-roosting bats are most commonly observed migrating in the autumn and only infrequently in the springtime (16, 17). Autumn flocks of eastern red bats were historically so notable that they would reportedly remain visible overhead for days at a time (18, 19). Because tree-roosting species likely have sex-partitioned summer habitat (16, 17, 20) and mate in the autumn (with delayed fertilization until parturition in spring and summer [20, 21]), it has been suggested that mating behavior may be a partial driver of autumn migratory patterns (22, 23). However, the patterns and drivers of migratory strategy in tree-roosting bats have rarely been examined quantitatively nor across their entire geographic range and annual cycles, and thus remain largely unknown.

Here we develop and apply a framework for understanding migration dynamics with a focus on North American tree-roosting bats, though our approach is amenable to other less-studied migratory animals (Figure 1). Our key methodological advance is integrating endogenous markers, in this case stable hydrogen isotope compositions of metabolically inert fur, into a hierarchical modeling framework that can reveal patterns of timing, distance, and direction of bat movement at a continental scale (Figure 1). Analysis of endogenous markers is a minimally-invasive and cost-efficient means of inferring location of tissue formation (24, 25). The basis for this approach is that stable hydrogen isotope compositions of precipitation vary across geographic and climatic gradients in broadly latitudinal bands (26). As tree-roosting bats molt during early-to mid-summer (27–29), their fur integrates isotopic compositions from the food and water they ingest from the local environment where the fur is synthesized. Therefore, stable hydrogen isotope compositions of bat fur reflect the summering grounds where fur was formed, even when sampled at other times (29, 30).

**Figure 1.**
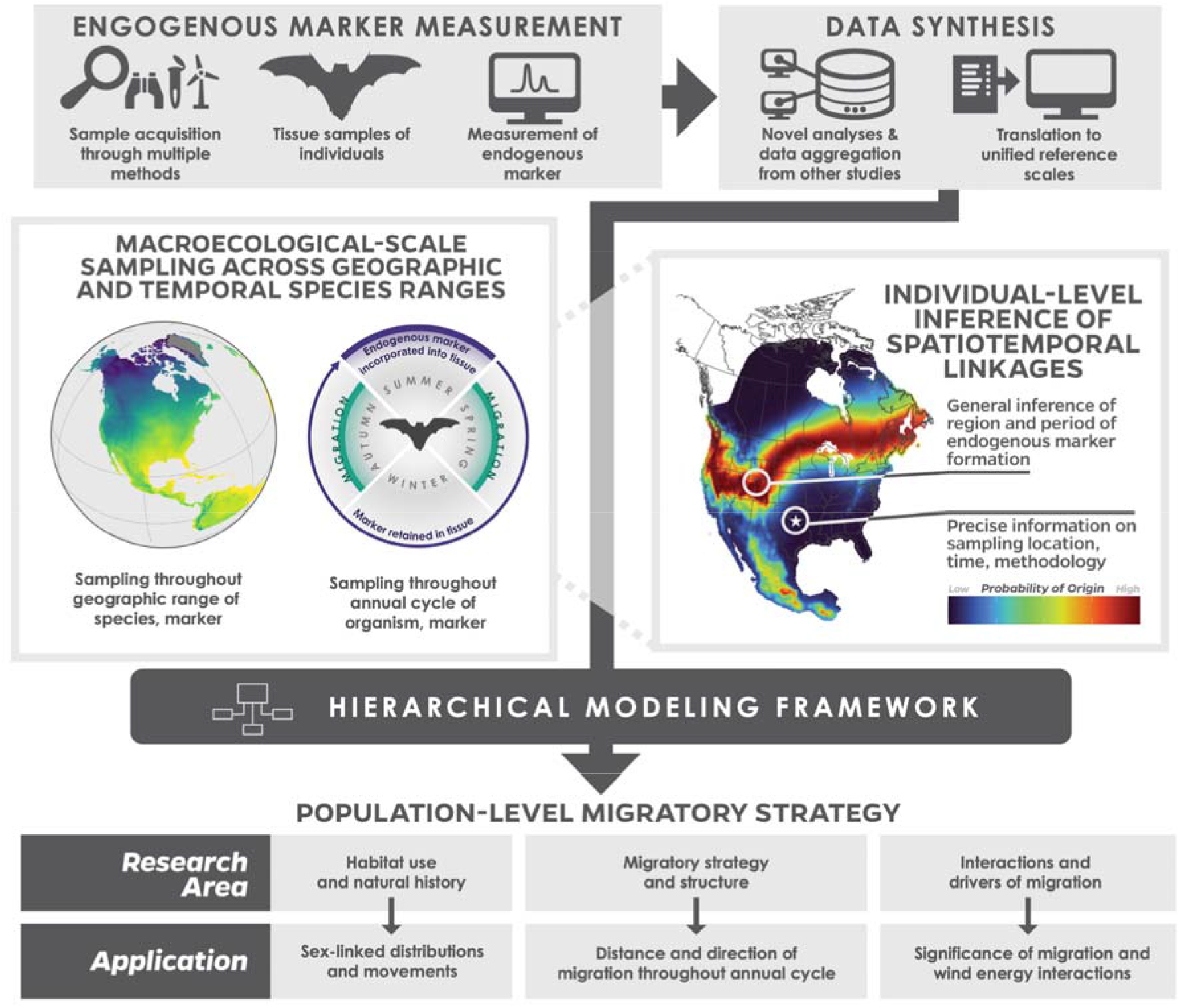
A workflow of the demonstrated research framework. At top, individual organisms were accessed through multiple methods, including live capture as part of monitoring studies, museum specimens, and carcass searches at wind energy facilities. Tissue samples were taken; in this case, of metabolically-inert keratin in hair, and analyzed for an endogenous marker for stable hydrogen isotope ratios. Next, data synthesis was conducted to assimilate the data collected for this study with previously-published data. To account for variation in methodologies and measurement standards from multiple laboratories, all data were transformed to a unified reference scale. Sampling was conducted across a broad geographic range, representing both the geographic ranges of the species but also broad sampling across the range of the marker (in this case, precipitation stable hydrogen isotope data, represented in the leftmost middle map with low values in purple [-200‰], median values in green [- 100‰], and high values in yellow [0‰]); it was also conducted across the annual cycle of the organism and its relationship with the marker. This widespread sampling enabled broad-scale individual-level inference of spatiotemporal linkages, where precise information on sampling location, time, and methodology could be linked with general inferences about the region and period of tissue formation represented by the endogenous marker, in this case the location and timing of summer molt. Both sources of data were incorporated into a hierarchical modeling framework, where hypotheses relevant to research areas including natural history, migration ecology, and behavioral ecology could be tested. Here, we applied these research areas to answer specific questions on sex-linked distributions, distance and direction of movement, and significance of migration and its interactions with bat fatalities at wind energy facilities.

Recent isotope-based studies suggest that tree-roosting bats can be long-distance migrants, although with high variation in distances traveled and unclear patterns of migration (27, 28, 31). However, prior studies had limited sample sizes and spatial extents (though see [32]). An advantage of our approach is the ability to synthesize large datasets comprising both new and published stable hydrogen isotope measurements transformed to a unified measurement scale. Our increased sampling across broad spatial extents and across as much of the annual cycle as possible (Figure 2) provides a means to integrate spatial information across two scales: precise spatial and geographic information reflecting where and when tissue was sampled and more general estimates of the time and location of tissue formation. Thus, our methodology accounts for variation inherent to sampling across multiple species, sampling locations and times, and analytical approaches.

**Figure 2.**
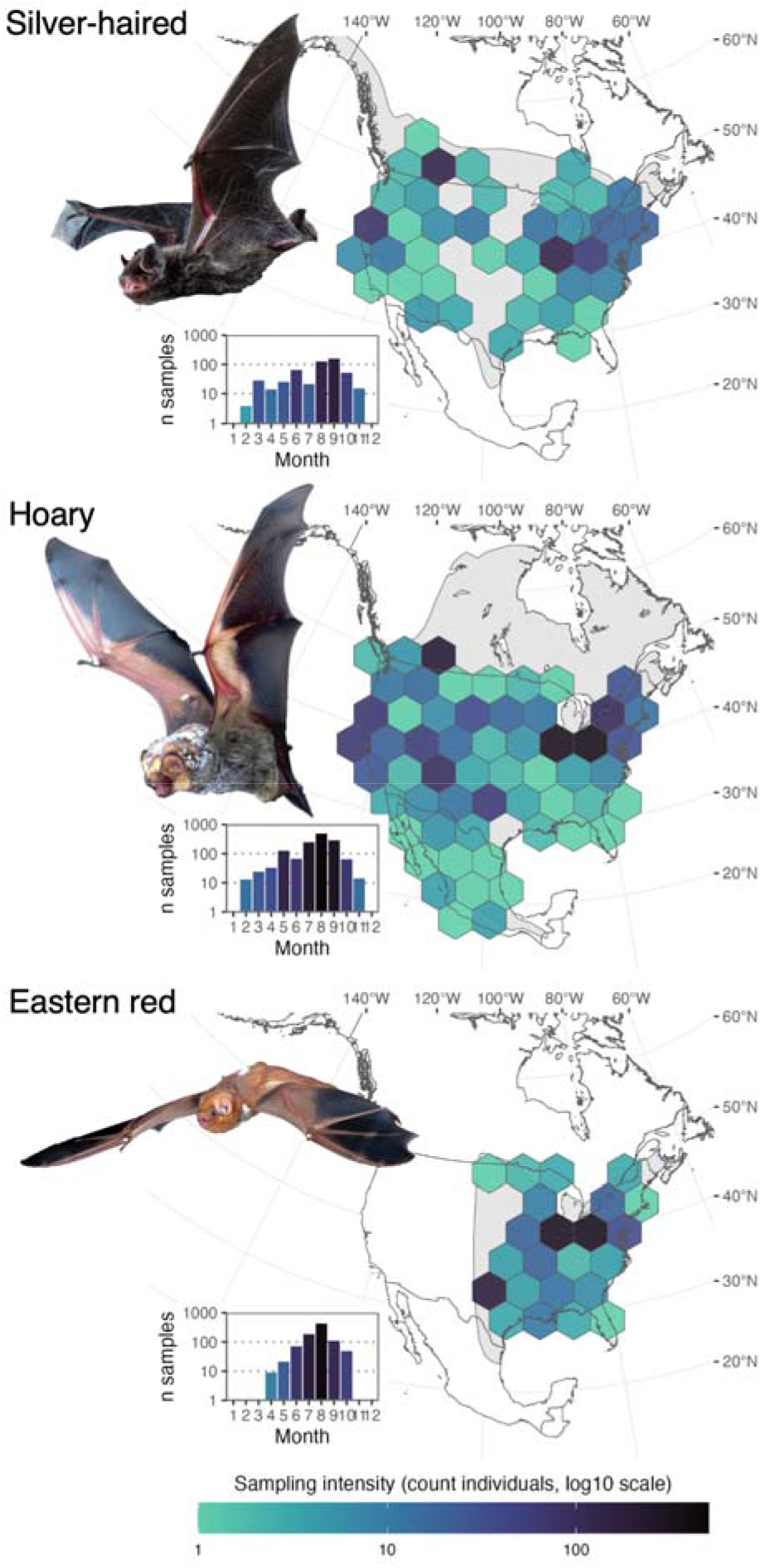
Geographic locations and month of collection of samples included in this study. Each panel depicts the sampling locations of samples binned within a 50x50km grid cell, overlaid above a gray-shaded species range map, for hoary, eastern red, and silver-haired bats respectively. Insets at bottom left of each panel show the number of fur samples obtained for each month of the calendar year (log10 scale). Photos by Sherri and Brock Fenton and José G. Martínez-Fonseca.

Better understanding of the migratory dynamics of tree-roosting bats has pressing conservation importance because these species face unprecedented large-scale mortalities at wind energy facilities. Collisions with the blades of wind turbines kill hundreds of thousands of bats in North America per year (33), with an as-yet-unexplained concentration of most fatalities among just a few species over just a few weeks of the year. Although some geographic variation exists across North America, a plurality (72-79%) of fatalities comprise just three species, hoary (*Lasiurus cinereus*; 32% (*Lasiurus borealis*; 24%), and silver-the haired bat (*Lasionycteris noctivagans*; 16%; 33, 34). These fatalities are concentrated in the late summer and early autumn (36, 37) and, given the time at which they occur, are circumstantially linked to migration (33, 36, 38, 39). However, the relationship between turbine impacts and migration remains largely speculative, and much research focused on mitigating impacts to bat species focuses on proximal attractive factors (e.g., movement by bats already nearby toward turbines; [38–40]). The rate of both existing and projected fatalities likely threatens the population viability of at least one species, the hoary bat (43, 44). As wind energy development is increasing rapidly as a key part of the transition to renewable energy, the urgency to explain and mitigate these interactions is growing (45).

We applied our methodology using measurements of endogenous markers from individuals to infer population-level migratory behaviors for the three bat species most impacted by wind energy development in North America. We sought to define the migratory strategies used by these species and discover how they relate to growing interactions with wind energy facilities. First, we tested for patterns of sex-biased distributions of summering grounds, expecting to find females would select higher-latitude summer habitat than males and that this factor may inform overall migratory strategy. We next examined patterns of distance and latitudinal direction of travel, expecting to find that bats that spent the summer at higher latitudes would travel furthest, and be more likely to move to lower latitudes to overwinter than counterparts that spent the summer at lower latitudes. We also expected to find evidence of migration to lower latitudes relative to summering grounds peaking in the winter months. Finally, we considered whether there were key differences in the migratory behaviors of bats sampled at wind energy facilities versus by other methods (e.g., live captures as part of monitoring studies and museum collection efforts). We control for key spatial, temporal, and geographic contexts, and thus unravel complex and counterintuitive signals of movement across the annual cycle of seasonal migrants. This application of individual-level data synthesis to quantify migration at the macroecological scale provides unparalleled insights into migratory strategies and potential risks to migratory organisms due to anthropogenic activities.

## Results

### Evidence of sex-biased latitudinal distributions of summer habitat

We hypothesized that female bats would be more likely to summer at higher latitudes than male bats, which we confirmed in all three species (mean estimate 0.67, 95% confidence interval [CI] 0.46-0.90; Table S3; Figure S3). Relative summering latitude of females was most marked in silver-haired bats, then hoary, and finally eastern red bats. The explanatory power (Tjur’s pseudo-R^2^) of the top model was 0.057.

### Unexpected patterns of movement for hoary and eastern red bats

We modeled the probability that individuals exhibited movement > 100 km and distance traveled of individuals with movement > 100 km using a two-part hurdle model, hypothesizing positive relationships between relative latitude of summer habitat and both likelihood of, and distance of, detected movement (Figure 3A). Silver-haired bats had a strong positive relationship between summer latitude and both likelihood of and distance of movement (Figure 3B-C); hoary bats had a weakly positive relationship of summer latitude with both variables; and eastern red bats had no apparent relationship between summer latitude and likelihood of travel and a weakly positive relationship between summer habitat and distance traveled when movement was detected (Figure 3B-C). We hypothesized a negative relationship between sampling latitude and both likelihood of and distance of detected movement (Figure 3D). All species had a negative relationship between sampling latitude and likelihood of movement (Figure 3E). Silver-haired bats had a negative relationship between sampling latitude and distance traveled (Figure 3F) as expected, but hoary bats had a positive and eastern red bats no relationship (Figure 3F; Table S4).

**Figure 3.**
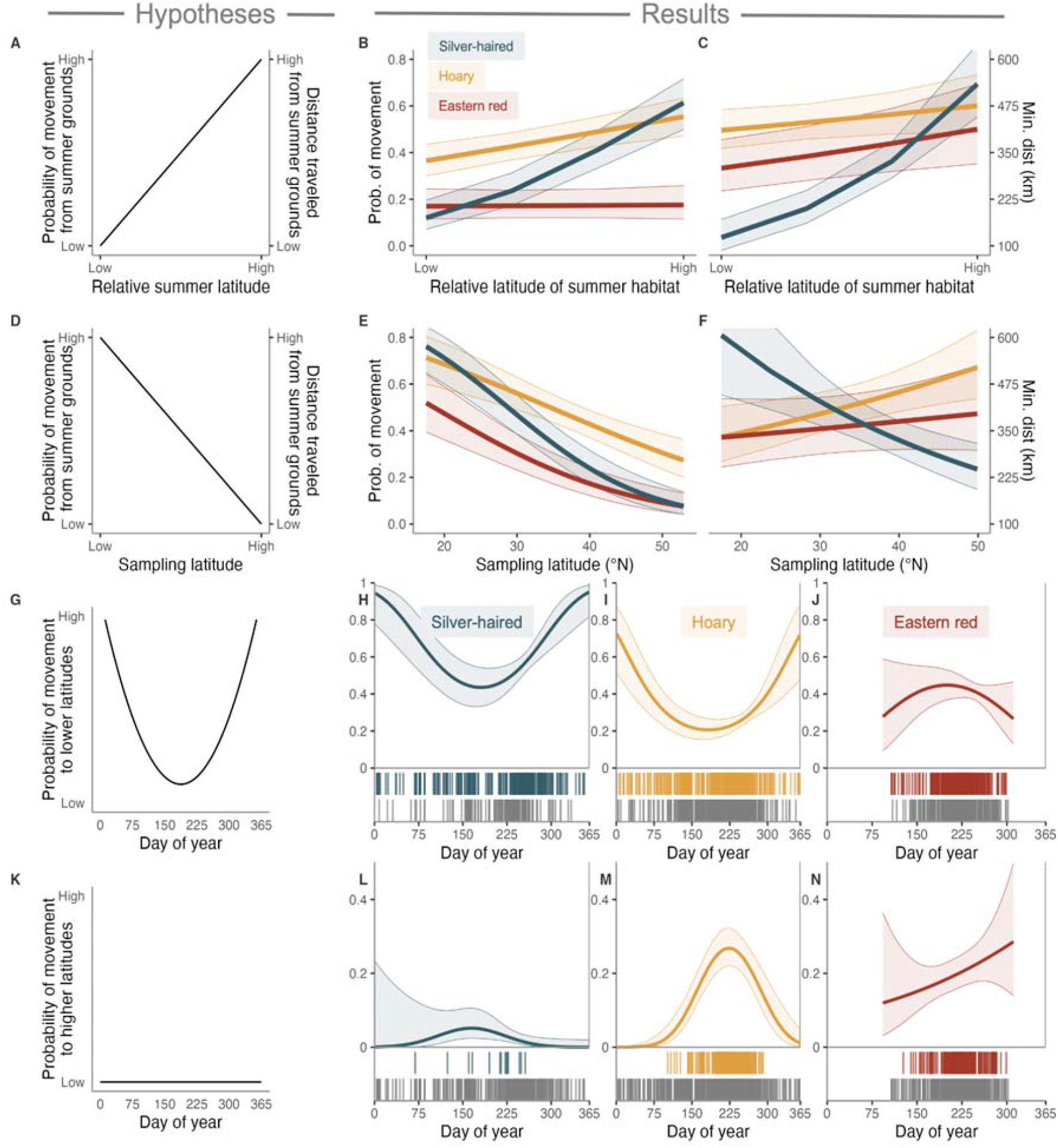
Hypothesized and realized results for distance and direction of travel by bats of each species. Top row: A positive relationship was predicted between the relative summer latitude of bats and both the probability of movement (left vertical axis) and distance of travel (right vertical axis, panel A). The probabilities of movement and distance traveled for silver-haired bats increased as expected, but with more ambiguous results for hoary and eastern red bats (panels B and C). We predicted sampling latitude was associated with a decrease in the probability of movement and distance traveled for all species (panel D), which was true for probability of movement for all species (panel E) but not for distance traveled by hoary and eastern red bats (panel F). Panels H-J and L-N show probability of movement in the indicated direction; colored carpet plots show number of individuals sampled with evidence of such movement; gray carpet plots show all individuals sampled without evidence of movement in that direction. We expected to find a U-shaped curve showing movement towards lower latitudes relative to sample site representing movements in between autumn and spring (panel G), which we broadly saw in all species (panels H-J), although with ambiguous results for eastern red bats that weren’t sampled throughout the entire annual cycle (panel J). We expected to discover little movement towards higher latitudes relative to sample sites (panel K), as in silver-haired bats (panel L), but we found substantial evidence of such movements in hoary and eastern red bats (panels M and N).

In general, hoary bats had the highest probability of movement and greatest minimum distance traveled when movement was detected (Figure S4A-B). Eastern red bats were least likely to have movement >100 km detected, but when moving >100 km they tended to travel farther than silver-haired bats (Figure S4A-B). Eastern red bats collected at wind energy facilities were more likely to have moved >100 km than those sampled by other methods (e.g., live capture; mean estimate 1.1, 95% CI [0.55, 1.7] ; Figure S4C). Silver-haired and hoary bats sampled at wind energy facilities were more likely to have traveled further than those sampled by other methods (Table S4; Figure S4D). Both model steps predicted a peak in likelihood of movement and distance traveled in the winter and spring relative to the summer (Figure S4A-B). There was no difference in probability of movement for bats of all species sampled through wind energy carcass surveys (mean estimate 0.07, 95%CI [-0.36, 0.51], but bats were associated with longer distances traveled when movement was detected (mean estimate 0.20, 95% CI [0.05, 0.43]). The two steps in the hurdle model had explanatory power of 0.094 (Thur’s pseudo-R^2^) and 0.349 (Nagelkerke’s pseudo-R^2^) respectively.

### Direction analyses indicate widespread movements to higher and lower latitudes

We hypothesized that movement to lower latitudes relative to summering grounds would be common, and greatest in the winter and least in the summer (i.e., form a U-shaped curve; Figure 3G). Conversely, we hypothesized that movement to higher latitudes relative to summering grounds would be rare and have no clear relationship with day of year (Figure 3K). We found that the probability of movement to lower latitudes was common in all species, and that movement to higher latitudes is common in the late summer/early autumn in hoary and eastern red bats (Figure 3H-J and L-N). Probability of movement to lower latitudes peaked during the winter months with probabilities > 0.72 and 0.95 for the two species with samples representing most of the year, hoary and silver-haired bats, respectively. Movement to higher latitudes was predicted with much higher probabilities for hoary and eastern red bats (peak probabilities of 0.28 and 0.45, respectively) than for silver-haired bats (peak predicted probability of 0.05). The probability of movement to higher latitudes for hoary bats was strongly unimodal, with the top 10% of predicted probabilities of movement in that direction occurring between days 209–237 (late July–late August). During this interval the probability of movement to higher latitudes was similar to that of to lower latitudes, with movement to higher latitudes having a slightly higher predicted probability (Figure 3H-J and L-N). Individuals that were sampled at wind energy facilities were more likely to have moved to higher latitudes than those sampled by other methods (mean estimate 0.29; 95% CI [.04, 0.54]; Table S5). The explanatory power (Tjur’s pseudo-R^2^) of the movement to lower latitudes model was 0.10 and 0.08 for the movement to higher latitudes model.

## Discussion

Migration drives broad changes to biodiversity and shifts ecosystem dynamics worldwide, but our knowledge of migration patterns and drivers for most taxa is limited. Here, we leverage new techniques to scale up information of individual-level movement inferred from synthesized stable isotope measurements to model range-wide patterns of migration in three species of tree-roosting bats. Our key finding is a novel migration pattern in which hoary and eastern red bats move counter to the expected direction towards lower latitudes in the autumn, and instead travel to higher latitudes from their summer habitats before ultimately reversing course to return to similar or lower latitudes to overwinter. This strategy provides critical insight into the behavioral drivers of the dramatic taxonomic and temporal concentrations of bat fatalities at wind energy facilities in North America, which has until now remained an ecological mystery. As we discuss below, our research framework provides a basis for inferring broad ecological- and conservation-relevant conclusions through incorporating rich endogenous marker data sampled across the spatial and temporal contexts of multiple species.

### Discovery of a pell-mell migratory strategy

Although most long-distance animal migrations comprise individuals tracking seasonal shifts in temperature or resource availability (7), our results show that hoary and likely eastern red bats move widely and long distances to higher and lower latitudes through late summer and early autumn, an unusual migratory pattern likely motivated by other factors. In hoary bats, which were widely sampled throughout the annual cycle in our study, these movement patterns are subsumed later in the autumn by an overall pattern of movement to lower latitudes, indicating that this species does generally migrate to more temperate regions relative to summering grounds. Migration of hoary bats in autumn has been described as “wandering” or circuitous, although such interpretations are based on tracking very few individuals during a few weeks (n=1–3; [12, 13]). Our dataset comprising thousands of individuals provides context for interpreting these tracking data: that is, there are three distinct range-wide migratory patterns in hoary and eastern red bats: 1) a “pell-mell” period, which we define here as widespread mixing across geographic regions in multiple directions, in late summer and early autumn; 2) a later (or perhaps overlapping) autumn migration to overwinter at generally lower latitudes; and 3) a spring migration towards higher-latitude summering grounds. Our confidence in this conclusion is supported by the contrast with silver-haired bat migration, which appear to follow a more conventional migratory strategy consistent with climate- or resource-seeking movements (patterns 2 and 3, above).

The pell-mell pattern of movement in autumn is more remarkable when considering that long-distance migration is both risky and typically a period of increased energetic demand. Most long-distance migrants employ strategies minimizing energetic expenditures and pursue efficient routes (46). The pell-mell movements speak to the potential capacity of hoary and red bats to migrate many hundreds to thousands of kilometers, much farther than strictly necessary to seek favorable climatic conditions or food availability (13, 47). Adaptive strategies, including the use of torpor (13, 48–50) and feeding during migration (51–53), may help them tolerate the energetic stresses of potentially traveling more than double the total distance between summer and winter habitats. We also note that partial migration—in which some individuals migrate but others do not—is likely common in these species (54, 55), as with migratory mammals generally (56).Because endogenous markers can be used to infer movement, but are only sensitive to movements across environmental marker gradients, we caution that use of stable isotopes may not be sensitive to the presence or degree of partial migration, and likely underestimate the proportion of migratory individuals generally. Our model results suggest that the vast majority of silver-haired and hoary bats likely overwinter at lower latitudes than where they spent the summer, although we were unable to infer this information for eastern red bats due to limited winter sampling (Figure 2). Evaluating the interactions of bioclimatic, social and behavioral, and physiological factors governing the capacities and decisions of individuals to migrate remains a promising future area of research (57, 58).

We found strong evidence of sex-differentiated summer distributions in all three species, a factor that might drive the multi-directional pell-mell autumn migration in hoary and red bats. Our results show that female bats were more likely than males to summer in high-latitude areas (Figure S3), supporting evidence of sex-biased migration that was previously inferred from distribution maps (20) and spatial variation in occurrence records (16, 17). All three species mate in the autumn, prior to or during autumn migration (21, 22). Although we found that summer distributions are biased by sex, autumn pell-mell movements appear to be undertaken by both sexes (see Supporting Results section). We expect that male bats may be incentivized to search for mating opportunities at higher latitudes than their summering grounds, either by seeking mates directly or traveling to hypothesized lekking or swarming locations (22, 23, 59). Female bats must also be incentivized to move in late summer, potentially to either seek out or avoid mating events, or possibly to pursue optimal prey after an interval of low mobility while raising young. An alternate driver of movements may be the incentive to seek out and track the mass emergence and movements of migratory insect prey (60, 61), which has been proposed as a driver of autumn movements to higher latitudes by migratory bats in Europe (62). Although such migratory events have been poorly characterized in North America, they could represent a spike in food availability in terms of thousands of tons of biomass (63).

### Migratory behavior as a driver of anthropogenic interactions

Our results provide the first direct support for the hypothesis that migration is a key driver of bat fatalities at wind energy facilities. To date, links of migratory behavior to wind energy impacts have been made circumstantially through the phenological overlap of autumn migration and peaks in bat fatalities at wind energy facilities (35, 38, 64). As migratory species may be more inclined to encounter or forage at the heights of wind turbines generally (60, 65), it has remained challenging to disentangle mechanisms driving fatalities from potentially spurious temporal correlations. Here, we demonstrate that the timing of the interval in which hoary and eastern red bats are regularly found both at both higher and lower latitudes relative to their summering grounds—likely representing an interval of widespread multi-directional movements by these species— coincides with the timing of fatalities. We found that the likelihood of movement to higher latitudes from summering grounds is greater among samples obtained at wind energy facilities, but not of the movement to lower latitudes which is more common later in the autumn. This further supports our conclusion that pell-mell migration is a key driver of bat fatalities at wind energy facilities.

Although substantial research has focused on proximate drivers of bat attraction to wind turbines (36, 40, 41, 64, 66, 67), a multi-directional “pell-mell” migration presents a significant ultimate driver of such interactions. If many hoary and eastern red bats move to higher latitudes relative to their summering grounds, and eventually move to lower latitudes to overwinter, the distances traveled by those migrants would be significantly greater than that of bats migrating in one direction to overwinter. Thus hoary and eastern red bats appear much more likely to encounter more obstacles during their seasonal movements. Simply put, their migratory behavior results in many more opportunities to encounter wind turbines than do species that do not engage in a long, likely wandering or circuitous, migration. This additional driver may help to explain why bat fatalities at U.S. and Canada wind energy facilities comprise predominately hoary (32%) and eastern red bats (24%), while other sympatric species with similar propensity for migration behavior– including the silver-haired and tricolored bat (*Perimyotis subflavus*) comprise a much smaller number (16% and 2%, respectively; [34]). Further, if individuals migrating through a given area are at greatest risk of fatally encountering wind turbines, that provides an explanation for why year-round monitoring of bat activity and modeling of habitat suitability are not strong predictors of bat fatality rates (68, 69).

### A framework for making broad inference from endogenous marker data

Here, we implemented a framework for unraveling migratory strategy and its interactions from the sampling of endogenous markers of individual organisms. Such approaches are growing in popularity, and include applications with stable isotopes, trace elements, genetics, and trait-based markers (70–76). By sampling with the greatest possible breadth with respect to the geographic range of the species, geographic variance of the marker, and annual cycle of both the species and the endogenous marker, we built a representative model using data from thousands of individuals to represent migratory strategy over the entire population and annual cycle of the species. We performed data synthesis which translated data collected at multiple laboratories to a unified reference scale. We also accounted for model error iteratively throughout the hierarchical modeling process, both within the origin mapping steps and at multiple stages when collecting summary statistics. This framework was then leveraged to provide insight into natural history and habitat use, migratory strategy and structure, and interactions and drivers of migration. We anticipate that this approach will be applicable to a diverse range of taxa across the Tree of Life.

## Materials and Methods

This study comprised data collection, synthesis, and analysis, including a hierarchical modeling framework to test key hypotheses (Figure 1). First, we collected and analyzed fur samples for stable hydrogen isotope compositions (□^2^H), and synthesized these data with those of other studies. We then modeled the relationships between □^2^H values of fur and local environment (precipitation) needed to infer individual origins and created individual-level models of probable origin of each fur sample. Next, we extracted key summary statistics about individual movements from those models, relying primarily on the geographic and temporal relationships between the region and time of probable fur origin and the location and timing of sampling. Finally, we used those summary statistics as predictors in a generalized linear modeling framework to identify drivers of sex-biased distributions, distance, and direction of travel.

### Sample and data collection

We collected samples of fur from between the scapula of 1,665 silver-haired (n=309), hoary (n=719), and eastern red bats (n=637) for analysis of □^2^H values. Most of these samples (84.7%) were obtained from carcasses salvaged at wind energy facilities, with the remainder coming from individuals that were captured alive. We integrated these new data with existing □^2^H values of fur for 1,286 individuals sampled primarily from live capture and museum specimens ([27–29, 77]; Table S1). In total, the dataset used in our study comprises □^2^H values from 545 silver-haired, 1,513 hoary, 893 eastern red bats representing broad geographic and intra-annual temporal extents (Figure 2). We compiled metadata associated with each individual, including species and morphology-based sex identification, geographic location of capture, and sampling method (e.g., carcass salvage, fur sampling of live individuals, or collection as a museum specimen). Geographic coordinates were compiled for each individual, either directly or through georeferencing locality descriptions.

### Assembly of new hydrogen stable isotope data

The 1,665 fur samples for which stable hydrogen isotope □^2^H analysis was performed in the present study were prepared as in (78). To account for exchange of keratin hydrogen with ambient vapor, we used comparative equilibration (79) in which samples were analyzed alongside matrix-matched international reference materials with known □^2^H values of non-exchangeable hydrogen and an internal standard. Fur samples were analyzed for □^2^H values using a ThermoFisher high temperature conversion/elemental analyzer (TC/EA) pyrolysis unit interfaced with a ThermoFisher Delta V+ (Thermo Fisher Scientific, Bremen, Germany) isotope ratio mass spectrometer. All preparation and analyses were conducted at the University of Maryland Center for Environmental Science Appalachian Laboratory.

### Assembly of stable hydrogen isotope data from previous studies

We assembled 1,286 □^2^H_fur_ measurements from prior studies. These isotopic compositions were generated using varying reference standards, many of which are laboratory-specific and in some cases not directly calibrated to the VSMOW-SLAP scale. To ensure data comparability, we used a previously described calibration chain transformation approach (80) to assure comparable □^2^H_fur_ values across all previous studies and our newly generated dataset (Figure S1). This process is based on the measured relationships of □^2^H values among secondary standards, which are then used to calibrate different reference standard scales (81). At a later model step, we account for uncertainty introduced by this transformation (next section).

### Relating fur and precipitation □^2^H values

The environmental drivers of spatial variation in □^2^H values of precipitation (□^2^H _precip_) are well understood, but ecological and physiological factors cause □^2^H_fur_ values to shift as hydrogen is incorporated into animal tissues. To model the relationship between bat fur and local precipitation □^2^H values, we relied on the measured □^2^H_fur_ values of bats sampled during the period of fur growth for each species, as defined by prior studies (28, 32, 77). We related each of these values to a modeled □^2^H value of precipitation at the sampling location, fitting a linear regression describing the predicted offset between □^2^H_precip_ and □^2^H_fur_ values (Table S2). The use of this regression enables prediction of the geographic region of origin of a sample based on its measured □^2^H_fur_ value and the modeled distributions of □^2^H_precip_ values.

### Probability-of-origin map creation and interpretation

We used a hierarchical framework to project the probability of origin for each sample of bat fur as described in (78). These assignments were conducted using the isotopeAssignmentModel tool in the R package isocat (78, 82). The output of this step is per-location probabilities of origin for each individual within each species’ geographic range, stored as a raster.

We next applied a clustering framework to identify groups of individuals sharing similar summer geographic origins to others of the same species. The general approach is to compare each probability-of-origin surface within a species to each other, determine their relative similarity, and group individual origins based on that similarity. Groupings of individuals were partitioned using k-means (Figure S2). The clustered origins of each group were arranged in ascending mean aggregate latitude to ordinally represent relative latitude of summer habitat compared to other origins from the same species.

Next, we calculated two key metrics to determine the characteristics of any detected movement by each individual: first, we measured the minimum distances traveled from summer origin to sampling location, and second the latitudinal direction (i.e., from higher or lower latitudes) of the origin with respect to sampling location. Each summary metric was designed to account for the uncertainty within geographic assignment of tissue origin, with distance traveled being calibrated by the performance of the metric for reference individuals and direction traveled using weighted subsampling to account for broad geographic areas of potential origins with similar likelihoods.

### Generalized linear modeling

We next used generalized linear models (GLM) to examine the relationship between demographic and geographic factors and the (1) sex identification, (2) minimum distance traveled, and (3) direction of travel of each individual. We developed separate models with these variables as responses, and included species, sampling day of year, relative latitude of summer origin, and sample source as wind-facility vs. non-wind-facility as interacting predictor variables. The first model, prediction of the sex identification, comprised a binomial regression to model sex (coded as 0 [male] and 1 [female]) on a dataset filtered to omit individuals for which sex was not reported (n = 677). The global model specification included the fixed effects: species, relative latitude of summer habitat, and sampling method; we also included the 2- and 3-way interactions of those predictors.

The response variable of the second model, minimum distance traveled, is zero-inflated and positive-skewed continuous data; therefore, we applied a two-part hurdle model (83). The first part was a binomial logistic regression of whether evidence of movement >100 km was present. The global model specification included the fixed effects: species, 2nd-degree polynomial of day of year (transformed to begin the year at day 150), relative latitude of summer habitat, sampling method, and latitude of sampling. Interacting effects were species with all other fixed effects, along with the interaction of day of year and relative latitude of summer habitat. The second part was a log-linked Gamma GLM using the same predictors as in the first to estimate the minimum distance traveled for individuals for which the minimum distance traveled was >100 km.

Our third model step twas fit to estimate the direction of travel of individuals. We were primarily interested in latitudinal movement (and most confident in our ability to infer this direction of travel given the broadly latitudinal nature of stable hydrogen isoscapes [25]), and summarized direction of travel to two probabilities: probability of movement to higher latitudes and of movement to lower latitudes. As these two probabilities likely should not sum to 1—as individuals may not have moved or moved detectably across isotopically distinct environmental conditions— we treated the likelihoods that an individual moved to higher or to lower latitudes as separate, and fit two binomial logistic regressions estimating the likelihood an individual had a detectable direction of movement. We fit these models to two subsetted datasets comprising individuals with evidence of movement to higher and to lower latitudes, respectively. For example, we fit a model predicting movement to lower latitudes using the fixed effects of species, day of year (beginning on day 150), and sampling method; we also included the interaction of species and day of year. We fit the model predicting movement to higher latitudes with the same predictors. We later conducted post-hoc Chi Squared tests of raw data to test for differences in sex ratios of reliably-identified individuals that moved to lower and to higher latitudes respectively.

All global models were fit to predictors and interactions selected for likely biological relevance. In cases where global models initially predicted complete separation, as in the direction-linked models which were limited to smaller sample sizes than the subsequent models, we iteratively eliminated predictors with respect to biological relevance until models no longer indicated complete separation. We iteratively dredged to compare among global models generated from all combinations of subsetted predictors. Models were ranked using an Akaike’s Information Criterion (AIC), and the model with the fewest degrees of freedom and ΔAIC < 2 was retained. If any variables contained variance inflation factors (VIF) ≥5, we removed the correlated variable with the lower coefficient. We then reran the model selection process until all VIF < 5 and the dredge function produced a stable top model. We checked the residuals of the linear regressions to ensure models did not significantly violate model assumptions and measured goodness-of-fit of our models to determine Tjur’s R^2^ for logistic regressions and Nagelkerke’s pseudo-R^2^ values for linear regressions.

## Acknowledgments

We would like to thank those involved in obtaining, sharing, and analyzing the samples used in this study; particular thanks to Robin Paulman, Regina Trott, Carlyle Meekins, Scott Osborn, Elizabeth Stevenson, J. Edward Gates, Cortney Pylant, Becky Johnson, Carl Herzog, Jason Williams, Anna Weaver, Greg Aldrich, J. Paul White, Dave S. Johnston, Charles Brown, Carly RW Kalina, Lisle Gibbs, and Bryan Carstens. We thank Michael Belitz, Ana Longo, Robert Fletcher, and Todd Katzner for their valuable feedback on the methodology and manuscript. This work was funded by the Maryland Department of Natural Resources and the U.S. Fish and Wildlife Service, as well as by a grant from the Competitive State Wildlife Grants Program to Ohio State University and the University of Maryland Center for Environmental Science as jointly administered by the US Fish and Wildlife Service, the Ohio Division of Wildlife and the Maryland Department of Natural Resources. CJC was supported by a University of Florida Graduate School Funding Award, a University of Florida Biodiversity Institute Fellowship, and Threadgill Dissertation Fellowship. This work was authored in part by the National Renewable Energy Laboratory, operated by Alliance for Sustainable Energy, LLC, for the U.S. Department of Energy Wind Energy Technologies Office. The findings and conclusions in this article are those of the author(s) and do not necessarily represent the views of the U.S. Fish and Wildlife Service, the Department of Energy, or the U.S. Government. The U.S. Government retains - and the publisher, by accepting the article for publication - acknowledges that the U.S. Government retains a nonexclusive, paid-up, irrevocable, worldwide license to publish or reproduce the published form of this work, or allow others to do so, for U.S. Government purposes.

