## Supplemental Materials for "Unusual migratory strategy a key factor driving interactions at wind energy facilities in at-risk bats"

### *Supporting Materials and Methods*

#### *Sample and data collection*

Coordinates of carcasses associated with wind-energy facilities were rounded to the nearest tenth of a decimal degree to obscure their association with specific facilities. When geographic coordinates were unavailable, the location was georeferenced to the county level using the geocode function of the ggmap R package (1) or matched with museum records in the GBIF database using the occ_search function of the rgbif R package (2).

#### *Assembly of new hydrogen stable isotope data*

The 1,665 fur samples for which hydrogen stable isotope analysis was performed in the present study were cleaned using 1:200 Triton X-100 detergent, 100% ethanol, and then air-dried in a fume hood (3). To account for exchange of keratin hydrogen with ambient vapor, we used comparative equilibration (4) in which samples were analyzed alongside matrix-matched international reference materials (USGS42, USGS43, CBS [Caribou Hoof Standard] and KHS [Kudu Horn Standard]; (3, 5)) with known $\delta^2$H values of non-exchangeable hydrogen (-72.9, -44.4, -157.0, and -35.3‰, respectively) and an internal standard (porcine keratin product #K3030; Spectrum Chemicals, New Brunswick, NJ, USA). Analytical precision of the internal keratin standard was 2.3‰ for $\delta^2$H. Approximately 0.15 mg of each cleaned sample and standard was weighed into silver capsules and exposed to ambient air for > 72 hrs prior to analysis for equilibration of exchangeable hydrogen. Fur samples were analyzed for $\delta^2$H values using a ThermoFisher high temperature conversion/elemental analyzer (TC/EA) pyrolysis unit interfaced with a ThermoFisher Delta V+ (Thermo Fisher Scientific, Bremen, Germany) isotope ratio mass spectrometer. Values of $\delta^2H_{fur}$ are reported in parts per mil (‰) on the Vienna Standard Mean Ocean Water-Standard Light Antarctic Precipitation (VSMOW-SLAP) scale. All preparation and analyses were conducted at the University of Maryland Center for Environmental Science Appalachian Laboratory.

#### *Relating fur and precipitation $\delta^2$H values*

Ideally, regressions between samples and environmental markers are fit to reference individuals of the same species sampled relatively evenly across isotopic and geographic space during the period of tissue formation (6). However, our dataset included uneven and temporally and geographically clustered sampling events. To account for challenges synthesizing data from different sampling regimes and geographic extents, species-specific variations in molt timing, and remaining variation across analysis laboratories and reference scales, we applied bootstrapped standardized major axis regressions to generate a suite of competing models relating modeled $\delta^2H_{precip}$ and the $\delta^2H_{fur}$ values of each species. We accounted for this sampling bias by: (1) iteratively sampling with replacement across 1,000 regressions where one sample was retained per sampling location (decimal coordinates) and sampling year, retaining models fit to ≥20 points; and (2) employing model filters including and excluding samples from wind facilities, which tended to be particularly clustered geographically. Additionally, we fit models to all individuals of each species and to a subset of those analyzed at a given laboratory.

We evaluated three different isoscape models of mapped $\delta^2H_{precip}$ values, representing annual and multi-month temporal extents (annual, IsoMAP job 66100 (7); June–August, IsoMAP job 66098; and growing season (8)). We selected top-performing models for each species based on the mean $R^2$ of the bootstrapped model, reporting mean coefficient estimates $\underline{\alpha_q}$ and $\underline{\beta_q}$ as intercept and slope for isoscape $q$, respectively.

To incorporate variation in instrument precision across analysis laboratories and reference standards, we calculated the residuals of each species- and laboratory-specific dataset to the top-performing model of that species ($\underline{\sigma_m}$, where *m* is the analysis laboratory). We included the standard deviation of the residuals with respect to each species and analysis laboratory (Table S2) in the subsequent probability-of-origin map creation step, directly propagating uncertainty there.

***Probability-of-origin map creation***

We used a hierarchical framework for projecting the probability of origin for each sample of bat fur as described in (9). Based on the best-fit bootstrapped major axis regression analysis corresponding with isoscape $q$ and mean coefficient estimates $\underline{\alpha_q}$ and $\underline{\beta_q}$ , the predicted $\delta^2H_{precip}$ value at summer origin for individual *i* ($X_i$) is:

$$X_i = \left(Y_{i,m} - \underline{\alpha_q}\right) / \underline{\beta_q}$$

where $Y_{i,m}$ is the measured $\delta^2H_{fur}$ value obtained from source *m*.

We calculated the probability of an individual's origin in a given location, as described in (10), using Bayes' Theorem to estimate the posterior probability of a normal distribution with a single sample:

$$f(X_i|\mu,\sigma) = \left(\frac{1}{\sqrt{2\pi\sigma^2}}\right) exp\left[-\frac{1}{2\sigma^2}(X_i - \mu)^2\right]$$

where $X_i$ is the predicted $\delta^2H_{precip}$ value at summer origin, μ is the isoscape value at the location, and $\sigma$ combined sources of error. Given $\sigma_{q,\mu}$ as the standard deviation at a given location in the isoscape model and $\underline{\sigma_m}$ as the mean of the residuals of known-origin individuals from laboratory *m*, error was combined as:

$$\sigma = \sqrt{\sigma_{q,\mu}{}^2 + \underline{\sigma_m}{}^2}$$

All isoscapes were projected into an equal area projection prior to analysis. We constrained the accessible area of assignment to a buffered region defined by the IUCN range maps for each species (11). The resulting surface was normalized to sum to 1. These assignments were conducted using the isotopeAssignmentModel tool in the R package *isocat* (9, 12). The output of this step is per-location probabilities of origin for each individual within each species' geographic range, stored as a raster.

***Probability-of-origin map interpretation***

We estimated the minimum distance each individual traveled between the nearest likely summer origin and its sample site by accounting for model performance on reference individuals and propagating probability-of-origin model error (9). To do so, we applied a nonparametric bootstrapping method. For each cell within an individuals' quantile probability of origin surface, we resampled the empirical distribution of reference quantile probabilities with replacement 100,000 times and calculated the proportion of times the cell value fell at or above the reference values. Reference values were generated from the quantile probabilities at sample sites for the reference individuals of the same species. The resulting surface, ranging from 0–1, thus reflects a level of certainty that any given potential origin resembles that of the origins of the reference individuals. We estimated the minimum distance traveled using a 75% likelihood threshold (surface value of 0.25), and calculated the minimum distance between all points above that threshold and the sampling site on an ellipsoid using the geosphere R package (13).

The general direction of movement between the molt location and the location at the time of sampling was estimated using a 10,000-fold resampling of potential origins. We subsampled each potential origin, weighted by the likelihood-of-origin of each raster cell multiplied by a conservatively-scaled distance-decay function, designed to weigh against origins excessively distant from the sample site (e.g., thousands of km). The angle of travel between each subsampled origin and the individual's sample location was recorded, and we calculated the proportion of subsampled points falling both at higher and lower latitudes relative to the sampling site. Individuals were classified as having moved to higher latitudes if a high proportion of selected points (>70%) from the summer origin had higher latitudes than the sampling location, and vice-versa for points at lower latitudes for migrants having moved to lower latitudes.

***Probability-of-origin map clustering***

As described by (9), we applied a metric of niche similarity (14) to evaluate the similarity of all probability-of-origin surfaces in a sequence of pairwise comparisons. We then populated a symmetric similarity matrix with niche similarity values, then applied hierarchical clustering using Ward's criterion (15, 16) to each distance matrix. Groupings of individuals were partitioned using k-means within-group sum-of-squares, with k groups of individuals evaluated at the elbow of the k vs. sum-of-squares line (Figure S3). The collective origins of each cluster of individuals were arranged in ascending mean aggregate latitude to ordinally represent relative latitude of summer origins to other origins represented in the same species.

***Generalized linear model selection***

All global models were fit to predictors and interactions selected for likely biological relevance. In cases where global models did not initially predict complete separation, as in the direction-linked models which were limited to smaller sample sizes than the subsequent models, we iteratively eliminated predictors with respect to biological relevance until models did not separate. We iteratively applied the dredge function from the R package *MuMIn* (17) to compare among global models generated from all combinations of subsetted predictors. Models were ranked using an Akaike's Information Criterion (AIC;(18)), and the model with the fewest degrees of freedom and $\Delta$AIC < 2 was retained. If any variables contained variance inflation factors (VIF) ≥ 5, we removed the correlated variable with the lower coefficient. We then reran the model selection process until all VIF < 5 and the dredge function produced a stable top model. We checked the residuals of the linear regressions to ensure models did not significantly violate model assumptions and measured goodness-of-fit of our models using the *performance* R package (19) to determine Tjur's $R^2$ for logistic regressions and Nagelkerke's pseudo-$R^2$ values for linear regressions.

***Generalized linear modeling of prediction of sex***

As the sex of carcasses obtained by wind-energy salvage can be misidentified, with a bias towards male identification more than half of the time (20–22), we only discuss sex-based model results from bats sourced using other methods, e.g., live capture.

***Quantitative analyses***

Statistical modeling was conducted in R version 4.1.2 (23). Beyond the packages specified above, we relied extensively on the tidyverse family of packages (24), isocat (9, 12), raster (25), and sf (26). Code used to run the analyses described in this study, as well as package version information, is available at https://github.com/cjcampbell/BatMigratoryDistDir and repositoried at https://doi.org/10.5281/zenodo.10578032.

### ***Supporting Results***

#### ***Generalized linear modeling of sex identification***

Uncertainty about the reliability of morphological sex identifications for most samples precluded using sex as a candidate predictor in the modeling framework (see above), but raw data indicated similar but unequal sex ratios among bats that moved to higher latitudes vs. those that did not for each species. Of bats reliably sexed (i.e., those with morphological identification based on samples not obtained at wind energy facilities), somewhat fewer female than male bats moved to higher latitudes, but with sex identifications ranging 43-59% female for each group. Pearson's Chi-squared tests did not show significant differences among the sex of bats that moved to higher latitudes vs. those that did not ($p = 0.40$ for silver-haired, 0.08 for hoary, and 0.07 for eastern red bats).

### ***References cited in supporting materials***


1. D. J. Kahle, H. Wickham, ggmap : Spatial Visualization with ggplot2. *R J.* **5**, 144 (2013).
2. S. Chamberlain, *et al.*, rgbif: Interface to the Global Biodiversity Information Facility API (2020).
3. T. B. Coplen, H. Qi, USGS42 and USGS43: Human-hair stable hydrogen and oxygen isotopic reference materials and analytical methods for forensic science and implications for published measurement results. *Forensic science international* **214**, 135–41 (2012).
4. L. Wassenaar, K. Hobson, Comparative equilibration and online technique for determination of non-exchangeable hydrogen of keratins for use in animal migration studies. *Isotopes in Environmental and Health Studies* **39**, 211–217 (2003).
5. L. I. Wassenaar, K. A. Hobson, Two new keratin standards ($\delta^2$H, $\delta^{18}$O) for daily laboratory use in wildlife and forensic isotopic studies. *Conference on Applications of Stable Isotopes* (2010) (April 5, 2016).
6. M. B. Wunder, D. R. Norris, "Analysis and Design for Isotope-Based Studies of Migratory Animals" in *Tracking Animal Migration with Stable Isotopes*, K. A. Hobson, L. I. Wassenaar, Eds. (2008), pp. 107–128.
7. G. J. Bowen, Z. Liu, H. B. Vander Zanden, L. Zhao, G. Takahashi, Geographic assignment with stable isotopes in IsoMAP. *Methods in Ecology and Evolution* **5**, 201–206 (2014).
8. G. J. Bowen, L. I. Wassenaar, K. a. Hobson, Global application of stable hydrogen and oxygen isotopes to wildlife forensics. *Oecologia* **143**, 337–348 (2005).
9. C. J. Campbell, M. C. Fitzpatrick, H. B. Vander Zanden, D. M. Nelson, Advancing interpretation of stable isotope assignment maps: comparing and summarizing origins of known-provenance migratory bats. *Animal Migration* **7**, 27–41 (2020).
10. M. B. Wunder, "Using isoscapes to model probability surfaces for determining geographic origins" in J. B. West, G. J. Bowen, T. E. Dawson, K. P. Tu, Eds. (Springer, 2010), pp. 251–270.
11. IUCN BSG, Chiroptera (spatial data) (2021) (September 14, 2021).
12. C. Campbell, isocat: Isotope Origin Clustering and Assignment Tools (2020).
13. R. J. Hijmans, geosphere: Spherical trigonometry (2019).
14. T. W. Schoener, Nonsynchronous spatial overlap of lizards in patchy habitats. *Ecology* **51**, 408–418 (1970).
15. J. H. Ward, Hierarchical grouping to optimize an objective function. *Journal of the American Statistical Association* **58**, 236–244 (1963).
16. F. Murtagh, P. Legendre, Ward's hierarchical agglomerative clustering method: which algorithms implement Ward's criterion? *J Classif* **31**, 274–295 (2014).
17. K. Bartoń, MuMIn: Multi-Model Inference (2020).
18. K. P. Burnham, D. R. Anderson, Model selection and multimodel inference: a practical information-theoretic approach, 2nd edn New York: Springer (2002).
19. D. Lüdecke, M. S. Ben-Shachar, I. Patil, P. Waggoner, D. Makowski, performance: An R package for assessment, comparison and testing of statistical models. *Journal of Open Source Software* **6** (2021).
20. J. M. Korstian, A. M. Hale, V. J. Bennett, D. A. Williams, Advances in sex determination in bats and its utility in wind-wildlife studies. *Molecular Ecology Resources* **13**, 776–780 (2013).
21. D. M. Nelson, *et al.*, Carcass age and searcher identity affect morphological assessment of sex of bats. *The Journal of Wildlife Management* **82**, 1582–1587 (2018).
22. A. S. Chipps, A. M. Hale, S. P. Weaver, D. A. Williams, Genetic approaches are necessary to accurately understand bat-wind turbine impacts. *Diversity* **12**, 236 (2020).
23. R Core Team, R: A language and environment for statistical computing. *R Core Team* (2016) https:/doi.org/3-900051-14-3.
24. H. Wickham, *et al.*, Welcome to the Tidyverse. *Journal of Open Source Software* **4**, 1686 (2019).
25. R. J. Hijmans, raster: Geographic data analysis and modeling (2020).
26. E. Pebesma, Simple features for r: Standardized support for spatial vector data. *The R Journal* **10**, 439–446 (2018).

***Supplemental Figures***

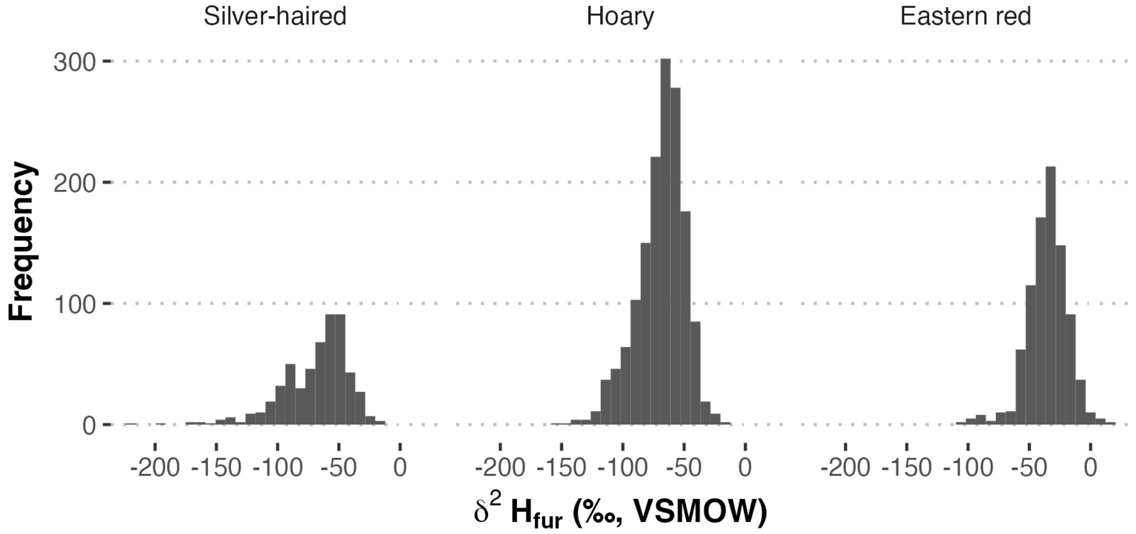


***Figure S1.*** *A histogram showing the frequencies of* $\delta^2H$ *measurements included in this study with respect to species, after synthesizing data measured across multiple reference scales to a single scale reflecting a unified reference standard (Vienna Standard Mean Ocean Water - Standard Light Antarctic Precipitation [VSMOW-SLAP]).*

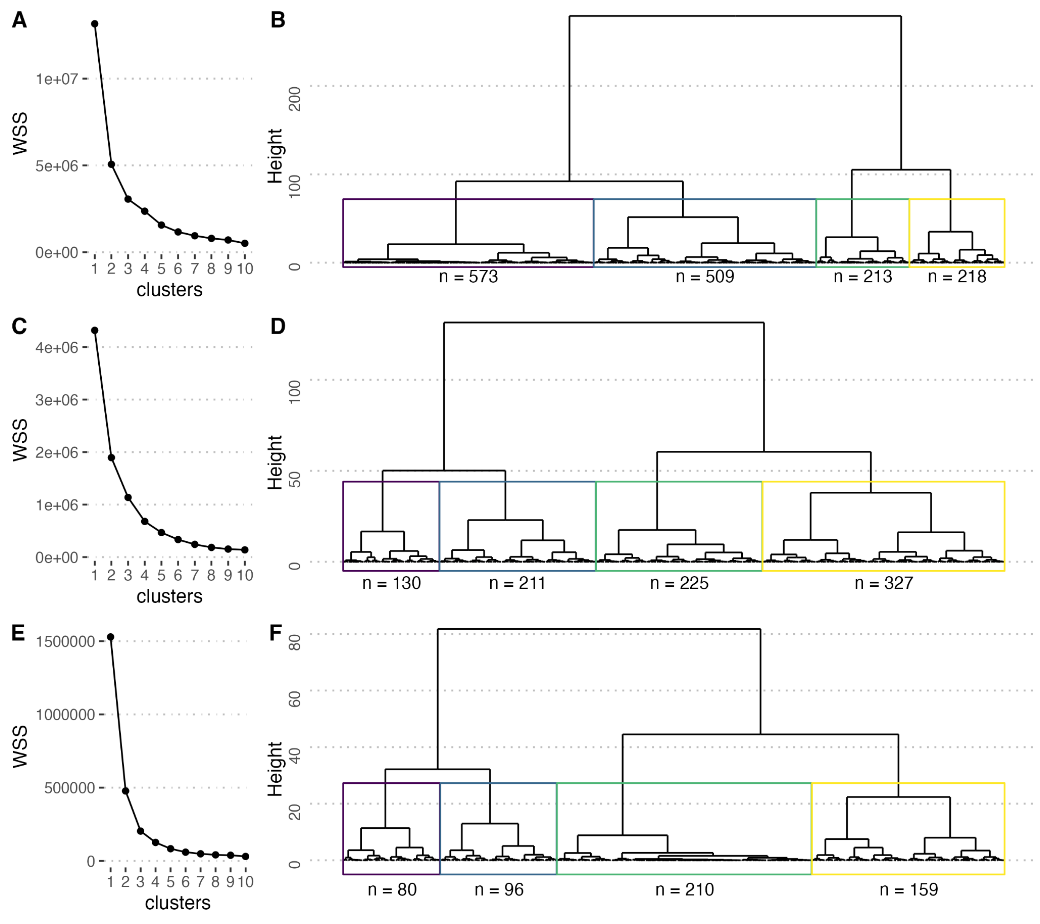


***Figure S2.*** *Results of clustering probability-of-origin surfaces and process of creating 'relative summer latitude' assignments. Left column shows k-means within-group sum of square (WSS) similarity-of-origin values for silver-haired (panel A), hoary (panel C), and eastern red (E), clustered into* k *groups. At right, dendrogram representations of the similarity of quantile probability-of-origin surfaces for silver-haired (B), hoary (D) and eastern red (F), grouped into k=4 groups of similar origins per species. Text at bottom indicates the number of individuals included in each group of similar summer origins per species.*

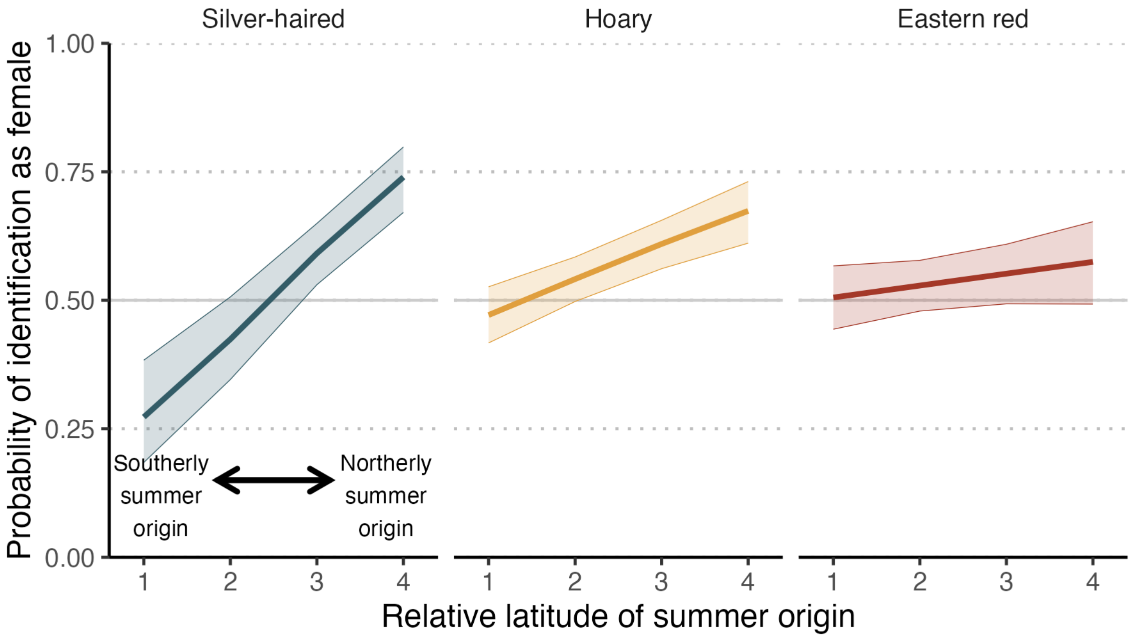


***Figure S3.*** *Model predictions for probability of sex as female. OriginCluster is a parameter summarizing relative likelihood of summer origin relative to other origins of the same species. All individuals were assigned to a relative latitude of summer origin group ranging from 1-4; a higher number represents a summer origin with higher relative latitude. Model predictions are projected for bats not sampled at wind turbines.*

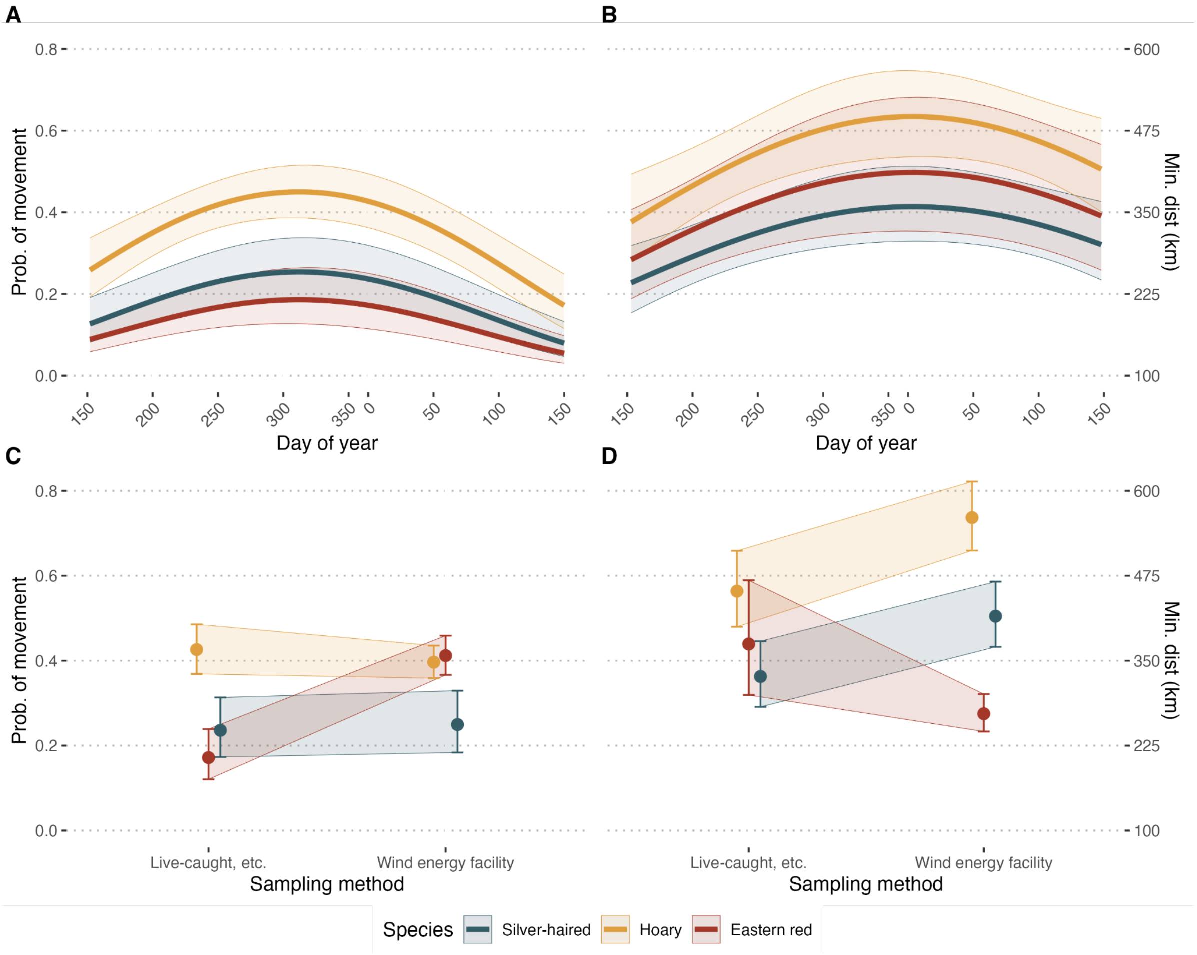


***Figure S4.*** *Model predictions for all interacting effects predicting probability of movement (left column) and minimum distance traveled (right column). Panels A and B show the effect of day of year, broken out by species. Panels C-D show the variation across species with respect to sampling method (was the sample taken from a carcass salvaged at a wind-energy facility, or by some other method).*

### ***Supplemental Tables***

***Table S1.*** *Number of individuals of each species analyzed for* $\delta^2H_{fur}$ *included in this study.*

| **Data source** | **Silver-haired** | **Hoary** | **Eastern red** | ***Total*** |
|---|---|---|---|---|
| This study | 309 | 719 | 637 | *1665* |
| Cryan *et al.* 2004 | 0 | 369 | 0 | *369* |
| Baerwald *et al.* 2014 | 119 | 176 | 0 | *295* |
| Pylant *et al.* 2014 | 0 | 0 | 112 | *112* |
| Pylant *et al.* 2016 | 0 | 249 | 144 | *393* |
| Fraser *et al.* 2017 | 117 | 0 | 0 | *117* |
| ***Total*** | *545* | *1513* | *893* | *2951* |

***Table S2.*** *A summary of top-performing bootstrapped transfer function model parameter estimates and selected isoscape model relating $\delta^2H_{fur}$ with isoscape-derived $\delta^2H_{precipitation}$ estimates. Testing individuals included in the model were individuals sampled during the literature-defined molt period (reported here in day of year) that were not sampled at wind energy facilities.*

| | | **Model parameter estimates** | | | | **Standard deviation of residuals for testing individuals** | | | | | |
|---|---|---|---|---|---|---|---|---|---|---|---|
| **Species** | **Isoscape** | **Intercept** | **Slope** | **$R^2$** | **Molt period (day of year)** | **This study** | **Cryan *et al.* 2004** | **Baerwald *et al.* 2014** | **Pylant *et al.* 2014** | **Pylant *et al.* 2016** | **Fraser *et al.* 2017** |
| Silver-haired | Annual | -3.80 | 0.64 | 0.64 | 171-238 | 16.5 | - | 21.4 | - | - | 8.0 |
| Hoary | Jun-Aug | -39.36 | 0.66 | 0.51 | 171-235 | 18.0 | 12.6 | 22.8 | - | 13.4 | - |
| Eastern red | Annual | 19.30 | 1.19 | 0.32 | 165-219 | 14.0 | - | - | 10.7 | 14.2 | - |

***Table S3.*** *Parameter estimate results for prediction of sex as female. The q-value, or false discovery rate correction for multiple testing, are bolded when values were less than the significance threshold q = 0.05, and parameter estimate cells are shaded blue for negative effect and red for positive effect in cases where q ≤ 0.05. The term "summer latitude" refers to an ordinal assignment of relative latitude of summer origin relative to other individuals of the same species.*

| | log(OR) | 95% CI | q |
|---|---|---|---|
| Intercept (Silver-haired) | -1.7 | -2.4, -0.97 | **<0.001** |
| Species | | | |
| *Hoary* | 1.3 | 0.50, 2.1 | **0.002** |
| *Eastern red* | 1.6 | 0.80, 2.4 | **<0.001** |
| Summer latitude | 0.67 | 0.46, 0.90 | **<0.001** |
| Wind-associated (no) | | | |
| *yes* | -0.8 | -0.99, -0.61 | **<0.001** |
| Species * summer latitude | | | |
| *Hoary * summer latitude* | -0.39 | -0.65, -0.15 | **0.002** |
| *Eastern red * summer latitude* | -0.58 | -0.84, -0.33 | **<0.001** |

***Table S4***. *Model results for two-step hurdle models evaluating evidence of movement > 100 km (step 1) and minimum distance traveled given movement > 100 km. Parameters are reported in log odds ratios (log(OR)) for the binomial logistic regression (step 1) and beta (step 2). The q-value, or false discovery rate correction for multiple testing, is bolded when less than significance threshold q = 0.05, and parameter estimate cells are shaded blue for negative effect and red for positive effect in cases where q ≤ 0.05. Parameter day of year is adjusted to begin on day 150 to center on early winter. The term "summer latitude" refers to an ordinal assignment of relative latitude of summer origin relative to other individuals of the same species.*

| | Step 1: evidence of movement | | | Step 2: Given movement, minimum distance traveled | | |
|---|---|---|---|---|---|---|
| | log(OR) | 95% CI | q | Beta | 95% CI | q |
| Intercept (Silver-haired) | 0.76 | -0.03, 1.6 | **0.071** | 5.4 | 4.9, 5.8 | **<0.001** |
| Day of year (centered on early winter) | | | | | | |
| *day of year* | -41 | -57, -27 | **<0.001** | -4.2 | -8.1, -0.05 | **0.038** |
| *day of year²* | -32 | -43, -20 | **<0.001** | -2.2 | -5.5, 1.1 | 0.2 |
| OriginCluster | 0.98 | 0.73, 1.3 | **<0.001** | 0.49 | 0.37, 0.62 | **<0.001** |
| Wind-associated (no) | | | | | | |
| *yes* | 0.07 | -0.36, 0.51 | 0.7 | .204 | 0.05, 0.43 | **0.015** |
| Sampling latitude | -0.1 | -0.13, -0.08 | **<0.001** | -0.03 | -0.04, -0.01 | **<0.001** |
| Summer latitude * species | | | | | | |
| *Summer latitude* Hoary* | -0.56 | -0.86, -0.28 | **<0.001** | -0.44 | -0.58, -0.31 | **<0.001** |
| *Summer latitude * Eastern red* | -0.81 | -1.1, -0.51 | **<0.001** | -0.39 | -0.54, -0.25 | **<0.001** |
| Wind-associated * species | | | | | | |
| *yes * Hoary* | -0.2 | -0.70, 0.31 | 0.5 | -0.03 | -0.26, 0.20 | 0.9 |
| *yes * Eastern red* | 1.1 | 0.55, 1.7 | **<0.001** | -0.56 | -0.85, -0.28 | **<0.001** |
| Sampling latitude * species | | | | | | |
| *sampling latitude * Hoary* | 0.05 | 0.03, 0.07 | **<0.001** | 0.04 | 0.03, 0.05 | **<0.001** |
| *sampling latitude * Eastern red* | 0.03 | 0.01, 0.05 | **0.016** | 0.03 | 0.02, 0.05 | **<0.001** |
| Day of year * Summer latitude | | | | | | |
| *day of year * Summer latitude* | 17 | 11, 22 | **<0.001** | 1.8 | 0.54, 3.1 | **0.005** |
| *day of year² * Summer latitude* | 9.1 | 4.5, 14 | **<0.001** | 0.09 | -1.0, 1.2 | 0.9 |

***Table S5.*** *Parameter estimate results for prediction of direction of travel. The q-value, or false discovery rate correction for multiple testing, are bolded when less than significance threshold q = 0.05, and parameter estimate cells are shaded blue for negative effect and red for positive effect in cases where q ≤ 0.05. Parameter day of year is adjusted to begin on day 150 to center on early winter.*

| | Movement to lower latitudes model | | | Movement to higher latitudes model | | |
|---|---|---|---|---|---|---|
| | log(OR) | 95% CI | *q* | log(OR) | 95% CI | *q* |
| Intercept (silver-haired) | 0.22 | 0.03, 0.40 | **0.039** | -3.8 | -4.5, -3.2 | **<0.001** |
| Species | | | | | | |
| *Hoary* | -1.2 | -1.4, -0.98 | **<0.001** | 2.3 | 1.7, 3.0 | **<0.001** |
| *Eastern red* | -0.52 | -0.78, -0.26 | **<0.001** | 2.4 | 1.8, 3.1 | **<0.001** |
| Day of year (centered on early winter) | | | | | | |
| *day of year* | 14 | 8.5, 20 | **<0.001** | | | |
| *day of year$^2$* | 5.7 | -3.2, 15 | 0.3 | | | |
| Wind-associated (no) | | | | | | |
| *yes* | | | | 0.29 | 0.04, 0.54 | **0.038** |
| Species * day of year | | | | | | |
| *Silver-haired * day of year* | 1.8 | -9.2, 12 | 0.8 | -18 | -46, 15 | 0.3 |
| *Hoary * day of year* | 0.46 | -7.7, 8.6 | >0.9 | 38 | 24, 53 | **<0.001** |
| *Eastern red * day of year* | 5.7 | -14, 25 | 0.7 | 14 | -6.2, 36 | 0.3 |
| *Silver-haired * day of year$^2$* | 24 | 12, 38 | **<0.001** | -42 | -99, -5.1 | 0.11 |
| *Hoary * day of year$^2$* | 18 | 9.8, 26 | **<0.001** | -44 | -62, -29 | **<0.001** |
| *Eastern red * day of year$^2$* | -17 | -38, 4.5 | 0.2 | 0.42 | -24, 23 | >0.9 |
